# A synthetic system that senses *Candida albicans* and inhibits virulence factors

**DOI:** 10.1101/342287

**Authors:** Michael Tscherner, Tobias W. Giessen, Laura Markey, Carol A. Kumamoto, Pamela A. Silver

## Abstract

Due to a limited set of antifungals available and problems in early diagnosis invasive fungal infections caused by *Candida* species are among the most common hospital-acquired infections with staggering mortality rates. Here, we describe an engineered system able to sense and respond to the fungal pathogen *Candida albicans*, the most common cause of candidemia. In doing so, we identified hydroxyphenylacetic acid (HPA) as a novel molecule secreted by *C. albicans*. Furthermore, we engineered *E. coli* to be able to sense HPA produced by *C. albicans*. Finally, we constructed a sense-and-respond system by coupling the *C. albicans* sensor to the production of an inhibitor of hypha formation thereby reducing filamentation, virulence factor expression and fungal-induced epithelial damage. This system could be used as a basis for the development of novel prophylactic approaches to prevent fungal infections.

## Introduction

Fungal pathogens cause diverse types of infections ranging from superficial to systemic and claim about 1.5 million lives per year^1^. *Candida* species are among the most common opportunistic fungal pathogens and represent the fourth-leading cause of hospital-acquired bloodstream infections with mortality rates of more than 40% in immunocompromised patients^1–4^.

*Candida albicans* represents the main cause of candidemia, being responsible for more than 40% of all cases worldwide^2^. *C. albicans* is a commensal colonizing the gastrointestinal and genitourinary tracts of healthy individuals^5^. However, under immunocompromised conditions, *C. albicans* can penetrate epithelia and cause severe systemic infections^6^. A key virulence factor of *C. albicans* is its morphologic plasticity; it can reversibly switch between a yeast and a filamentous or true hyphal morphology depending on the environmental conditions. On epithelial surfaces, initial adhesion and subsequent filamentation are required for efficient epithelial penetration^7^. In addition, several virulence factors such as Candidalysin, a peptide that damages epithelial cells, are produced exclusively by the hyphal form^8^.

Difficulties associated with rapid and reliable diagnosis as well as lack of specific disease manifestations contribute to the high mortality rates of systemic fungal infections^9^. Antifungal prophylaxis is currently applied to high risk individuals. While effective in preventing *Candida* infections^10,11^, prolonged prophylaxis can lead to the emergence of resistant strains or to an increase of species, that are intrinsically resistant to the limited set of antifungals available^10,12^. Development of novel antifungals with fungistatic or fungicidal activity is limited by the relative dearth of specific targets in these eukaryotic pathogens. Thus, more recently approaches targeting specific virulence factors have been gaining interest. This strategy can increase the number of potential targets, that are specific to the pathogen, and will exert less evolutionary pressure for the development of resistance compared to growth inhibition. In the case of *C. albicans*, inhibition of filamentation represents an important target because it could prevent physical penetration of epithelia as well as expression of hypha-specific virulence genes^7^.

A number of studies have shown that members of the human microbiota can inhibit hypha formation by *C. albicans*^13–16^. However, the inhibitory substance is usually produced constitutively and does not represent a specific response to the presence of the fungus. One example of a bacterial secreted molecule that efficiently blocks hypha formation by *C. albicans*, is cis-2-dodecenoic acid (BDSF), a diffusible signal factor that is produced by *Burkholderia cenocepacia*^13^.

Synthetic biology approaches using genetically engineered bacteria have demonstrated great potential in the prophylaxis or treatment of cancer, metabolic disorders and infectious diseases^17–20^. Using engineered microbes offers several advantages over traditional therapeutics including lower cost and higher specificity due to more targeted treatment^17^. A number of approaches using engineered bacteria to counteract or prevent bacterial and viral infections have been reported^21–26^. However, there have been no reports describing the use of reprogrammed microorganisms for treatment of or prophylaxis against fungal infections.

In this study, we developed an engineered system able to sense a fungal pathogen and respond by inhibition of fungal virulence factors. We discovered hydroxyphenylacetic acid (HPA) as a novel molecule secreted by *C. albicans* and used engineered *E. coli* cells carrying a HPA sensor to detect the presence of the fungus. Furthermore, we developed a sense-and-respond system by coupling the sensor to the production of BDSF to inhibit filamentation and virulence gene expression in *C. albicans* thereby protecting epithelial cells from fungal-mediated damage.

## Results

### System design

The overall goal of this work was to construct and characterize an engineered, commensal *E. coli* strain capable of first, detecting the presence of *C. albicans* and second, producing a hypha inhibitor in response to detection thereby preventing filamentation (**Figure 1**). The sensor should be able to sense a substance secreted by the fungus and control expression of a hypha inhibition module, which will lead to the production of an inhibitor of hypha formation. Thus, bacteria carrying this system would be able to protect an epithelium from being damaged by *C. albicans*.

**Figure 1.**
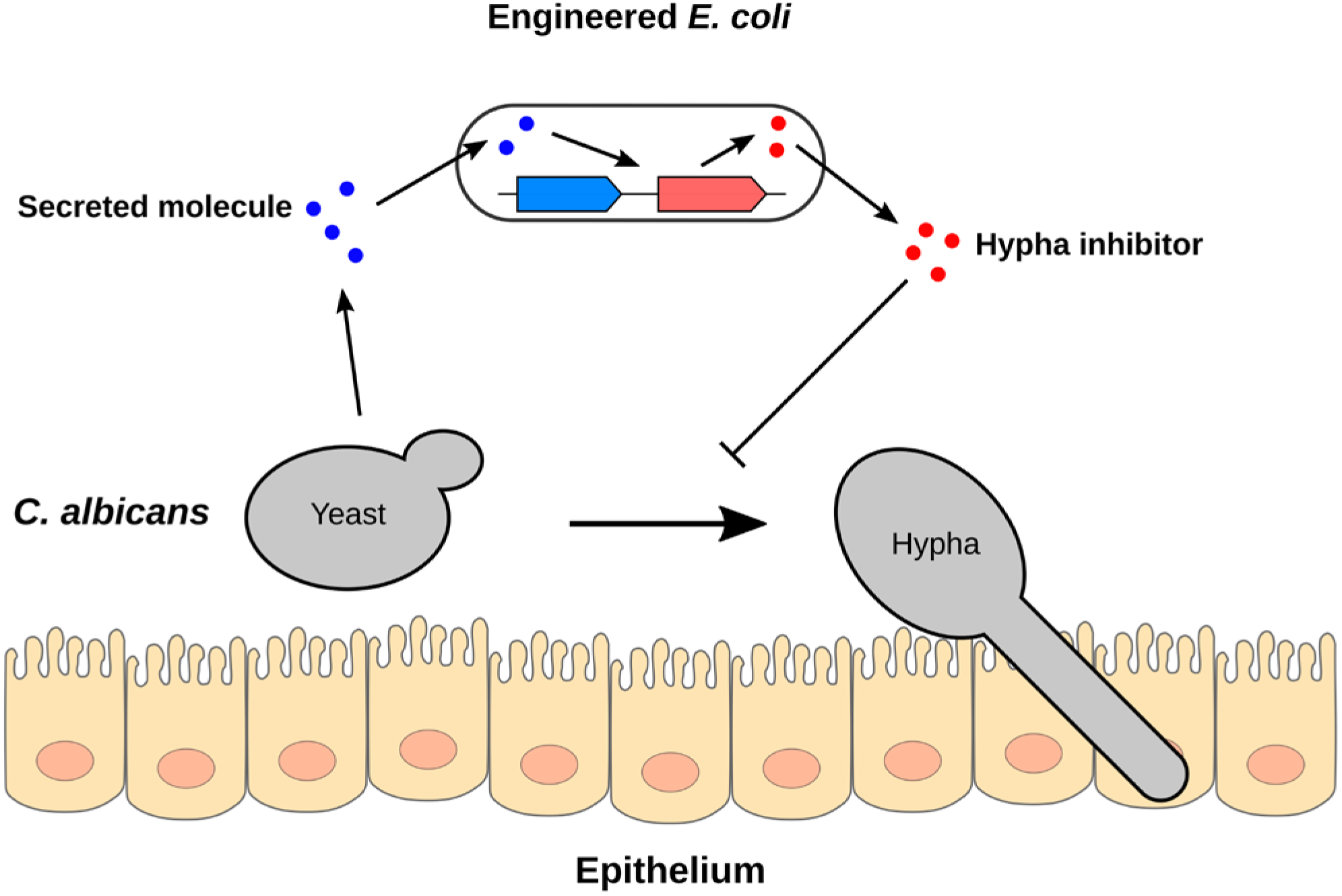
Design of a *C. albicans* sensing and hypha inhibition system. *C. albicans* can penetrate host epithelia by switching from a yeast to a hyphal morphology. Engineered *E. coli* cells can sense *C. albicans* secreted molecules and respond by production of an inhibitor of hypha formation thereby preventing epithelial damage and penetration.

### Engineered *E. coli* can sense a molecule produced by *C. albicans*

To engineer *E. coli* to sense C*. albicans*, we first focused our efforts on the known fungal quorum sensing molecule tyrosol (4-hydroxyphenylethanol), which is constitutively produced under various conditions including growth within a patient^27^. *E. coli* W strains are able to launch a transcriptional response to the structurally related substance 4-hydroxyphenylacetic acid (4-HPA)^28^. Therefore, we reasoned that the structural similarity between tyrosol and 4-HPA could serve as a starting point for the construction of a tyrosol sensor.

We first tested for tyrosol production by *C. albicans*. We harvested *C. albicans* culture supernatants and confirmed tyrosol production by HPLC-MS, which increased upon supplementation of the medium with its precursor tyrosine (**Figure S1a**). As a control we also determined the levels of HPA in fungal culture supernatants. Surprisingly, we detected HPA production by *C. albicans* and the levels increased upon tyrosine supplementation (**Figure 2a**). Furthermore, we could detect HPA in yeast and hyphal supernatants (**Figure S1b**). Based on its retention time we were able to identify the secreted molecule as being either 3-HPA or 4-HPA. Thus, *C. albicans* is able to produce HPA presumably via a pathway similar to tyrosol.

**Figure 2.**
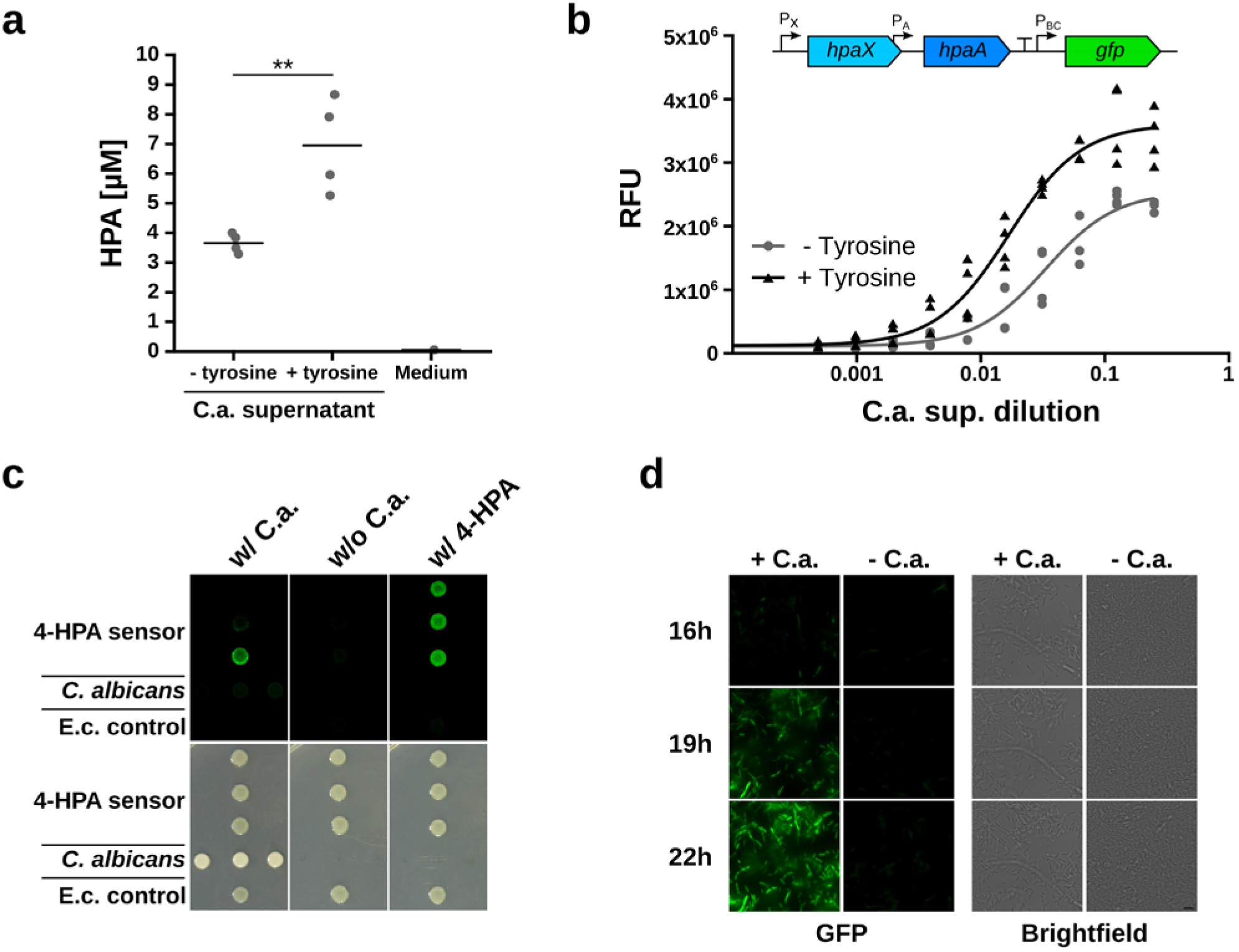
Construction of a *C. albicans* sensor. (a) *C. albicans* produces hydroxyphenylacetic acid (HPA). HPA levels in culture supernatants were determined by HPLC-MS. Addition of tyrosine increases HPA production. Means of biological replicates are indicated. One-tailed Student’s t-test: **P<0.01; (b) Dose-response of *E. coli* PAS691 to *C. albicans* culture supernatant. GFP was placed under the control of the 4-HPA inducible *P_BC_* promoter. HpaX imports 4-HPA into the cell and HpaA activates the *P_BC_* promoter upon binding to 4-HPA. A four-parameter log-logistic dose-response model was fitted to the data. (c) *E. coli* 4-HPA sensor (PAS691) senses *C. albicans*. *E. coli* and *C. albicans* were cocultured on LB agar plates supplemented with glycerol at 37°C. Sensor induction was determined after 24 hours of coculture by imaging using GFP excitation/emission filters (upper panel) and brightfield (lower panel). E. c. control cells carried the empty plasmid (PAS697). 4-HPA was added to the medium as indicated. (d) Time-lapse microscopy of *E. coli* PAS691 and *C. albicans* coculture in liquid minimum medium at 37°C. Images were taken at the indicated time points. Scale bar represents 5 μm.

*E. coli* has the ability to respond to 4-HPA and can be engineered to record its production by *C. albicans*. The *E. coli* 4-HPA sensor found in W strains consists of a transporter (HpaX) and a transcription factor (HpaA)^28,29^. Upon import by HpaX and binding to HpaA, 4-HPA can activate transcription from the downstream *P_BC_* promoter^28^. To test this system, we constructed a sensor plasmid carrying the *hpaX* and *hpaA* genes and the *P_BC_* promoter upstream of a *gfp* encoding gene (**Figure 2b**). Cells carrying the sensor plasmid (PAS691) showed increased GFP expression upon induction with 4-HPA (**Figure S1c**). Furthermore, the absence of the HpaX transporter (PAS692) decreased the sensitivity of the sensor. Tyrosol did not lead to efficient induction even in the presence of HpaX. To determine if the 4-HPA sensor can detect *C. albicans*, we incubated *E. coli* PAS691 with different dilutions of fungal supernatants and measured GFP levels after 24 hours (**Figure 2b**). *C. albicans* culture supernatants were able to strongly induce the 4-HPA sensor and tyrosine supplementation of the fungal culture further increased the induction level.

The sensor *E. coli* can directly detect the presence of *C. albicans*. We spotted *C. albicans* on agar plates and incubated them for 24 hours to allow for the production of HPA. Then, PAS691 sensor cells were spotted in proximity onto the same agar plate and incubated for additional 24 hours prior to imaging. Only bacterial cells in close proximity to *C. albicans* showed an elevated GFP signal, which declined with increasing distance from the fungal cells consistent with induction by a diffusible compound produced by *C. albicans* (**Figure 2c**). Importantly, we observed no significant induction in the absence of *C. albicans* and increased fluorescence upon addition of 4-HPA to the medium. We observed variation in the GFP intensity within PAS691 sensor spots, which might be due to copy number variation of the sensor-encoding plasmid. *C. albicans* spots showed low levels of autofluorescence. Furthermore, we also incubated *E. coli* PAS691 in liquid medium in the absence or presence of *C. albicans* and observed GFP induction only when the fungus was present (**Figure 2d**). Thus, our *E. coli* 4-HPA sensor can be used to detect *C. albicans*.

### Engineered *E. coli* can inhibit filamentation

*E. coli* can be engineered to induce production of BDSF, an inhibitor of *C. albicans* hypha formation that is produced naturally by *B. cenocepacia*^13^. The enzyme responsible for BDSF production has been identified as *B. cenocepacia* RpfF and expression in *E. coli* leads to production and secretion of BDSF^30^. To enable inducible RpfF expression in *E. coli* we placed the *rpfF* gene downstream of an anhydrotetracyline-(aTC-)inducible promoter (**Figure 3a**). RpfF has dehydratase and thioesterase activities and is able to produce BDSF from the acyl carrier protein (ACP) thioester of 3-hydroxydodecanoic acid, a fatty acid synthesis intermediate^30^.

**Figure 3.**
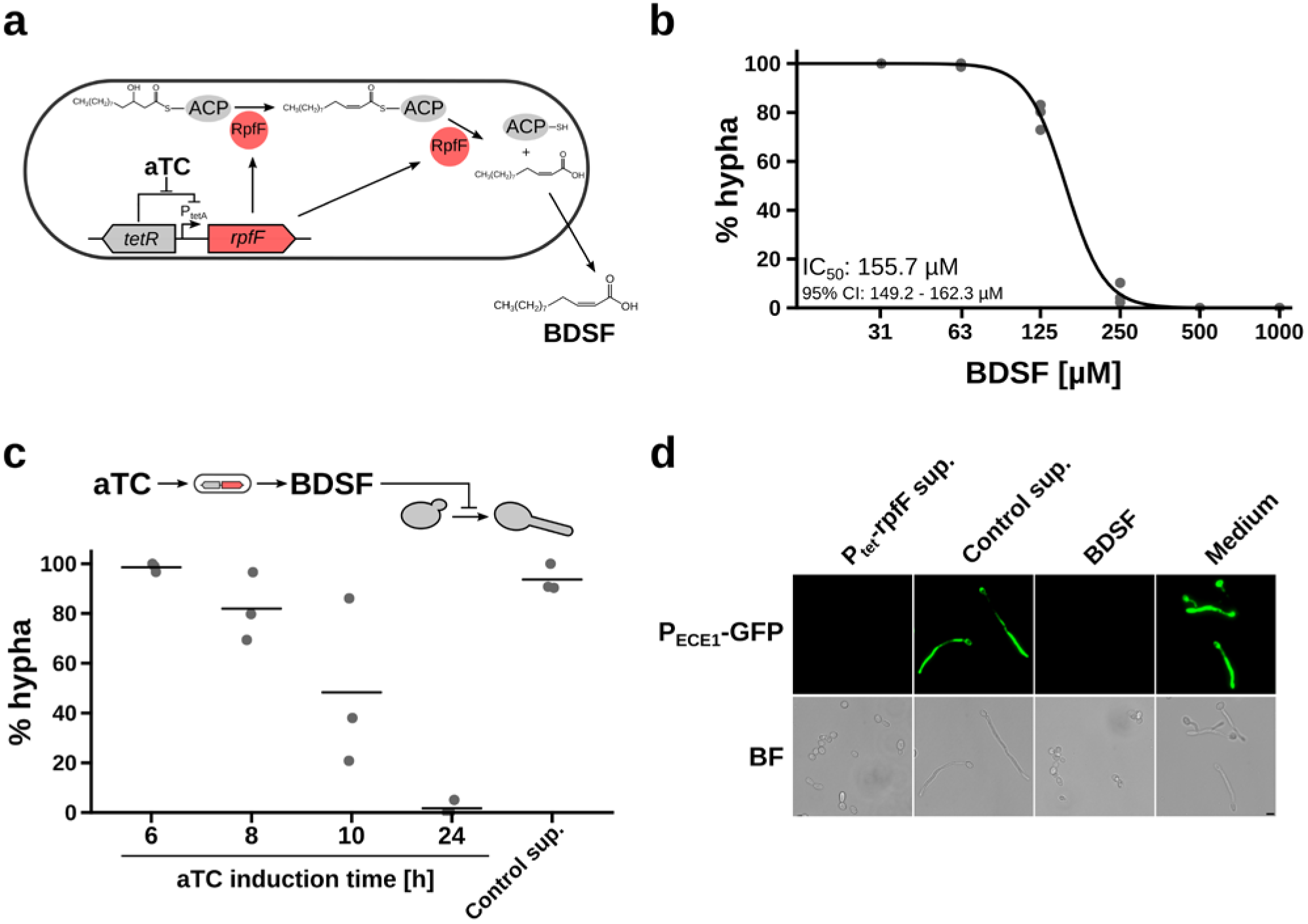
Filament inhibition by engineered *E. coli*. (a) Anhydrotetracycline‐ (aTC-)inducible system for the production of BDSF (cis-2-dodecenoic acid). Addition of aTC induces expression of RpfF, which converts 3-hydroxydodecanoic acid-acyl carrier protein (ACP) to BDSF. (b) Filament inhibition by BDSF. *C. albicans* yeast cells were treated with BDSF under hypha inducing conditions in DMEM medium with 5% FBS at 37°C and morphology was determined after 3 hours. A four-parameter log-logistic dose-response model was fitted to the data. 95% CI: 95% confidence interval; (c) *C. albicans* yeast cells were treated with supernatants from RpfF expressing *E. coli* cells (PAS689) under hypha inducing conditions in DMEM with 5% FBS at 37°C and morphology was determined after 3 hours. Control supernatants were obtained from MG1655 cultures. Means of biological replicates are indicated. (d) *C. albicans* cells containing an *ECE1* promoter-GFP reporter (CA58) were treated as described in (c) with *E. coli* supernatants after 24 hours aTC induction or with 1 mM BDSF. Scale bar represents 5 μm.

First, we confirmed that *C. albicans* hypha formation can be inhibited by BDSF (**Figure 3b**). To test our BDSF producing cells, we induced bacterial cells with aTC, harvested culture supernatants at different time points and used them for hypha inhibition assays. Starting at 8 hours post induction *E. coli* supernatants were able to inhibit *C. albicans* filamentation (**Figure 3c**). Thus, *E. coli* cells carrying the aTC-inducible system (PAS689) are able to produce BDSF at a level, that is sufficient to inhibit *C. albicans* hypha formation.

BDSF produced by *E. coli* is able to inhibit virulence gene expression in *C. albicans*. We used a fungal reporter strain containing GFP fused to the promoter of the *ECE1* gene (CA58) encoding the precursor of Candidalysin^8^. We determined GFP expression levels upon hypha induction with fetal bovine serum (FBS) at 37°C and observed strong *ECE1* induction (**Figure 3d**). However, upon treatment with BDSF or aTC-induced *E. coli* supernatants we observed no GFP induction. These data indicate that *E. coli* cells producing BDSF can inhibit hypha formation and virulence gene expression in *C. albicans*.

### Protection of epithelial cells by engineered bacteria

BDSF protects epithelial cells from killing by *C. albicans*. We incubated monolayer of Caco-2 epithelial cells with *C. albicans* in the absence or presence of BDSF. Without inhibitor fungal cells start to filament extensively in the coculture, whereas in the presence of BDSF only yeast cells were observed after 9 hours and 24 hours interaction, respectively (**Figure 4a**). To determine epithelial cell damage we measured lactate dehydrogenase (LDH) release in mammalian-fungal cocultures treated with increasing concentrations of BDSF. As expected, we observed a dose-dependent decrease in *C. albicans*-induced cytotoxicity with increasing BDSF concentrations (**Figure 4b**). Furthermore, the concentration required for epithelial cell protection matches the concentration needed for hypha inhibition (**Figure 3b**). Interestingly, we observed a similar protective effect of BDSF when we cultured epithelial cells in transwell plates and treated them with *C. albicans*. The decrease in transepithelial electrical resistance (TEER) caused by *C. albicans* was markedly reduced in the presence of 500 μM BDSF (**Figure S1d**). In addition, the increase in epithelial permeability upon treatment with fungal cells was also reduced in the presence of BDSF (**Figure S1e**). Furthermore, BDSF alone did not show any cytotoxicity (**Figure 4b** and **Figure S1d**).

**Figure 4.**
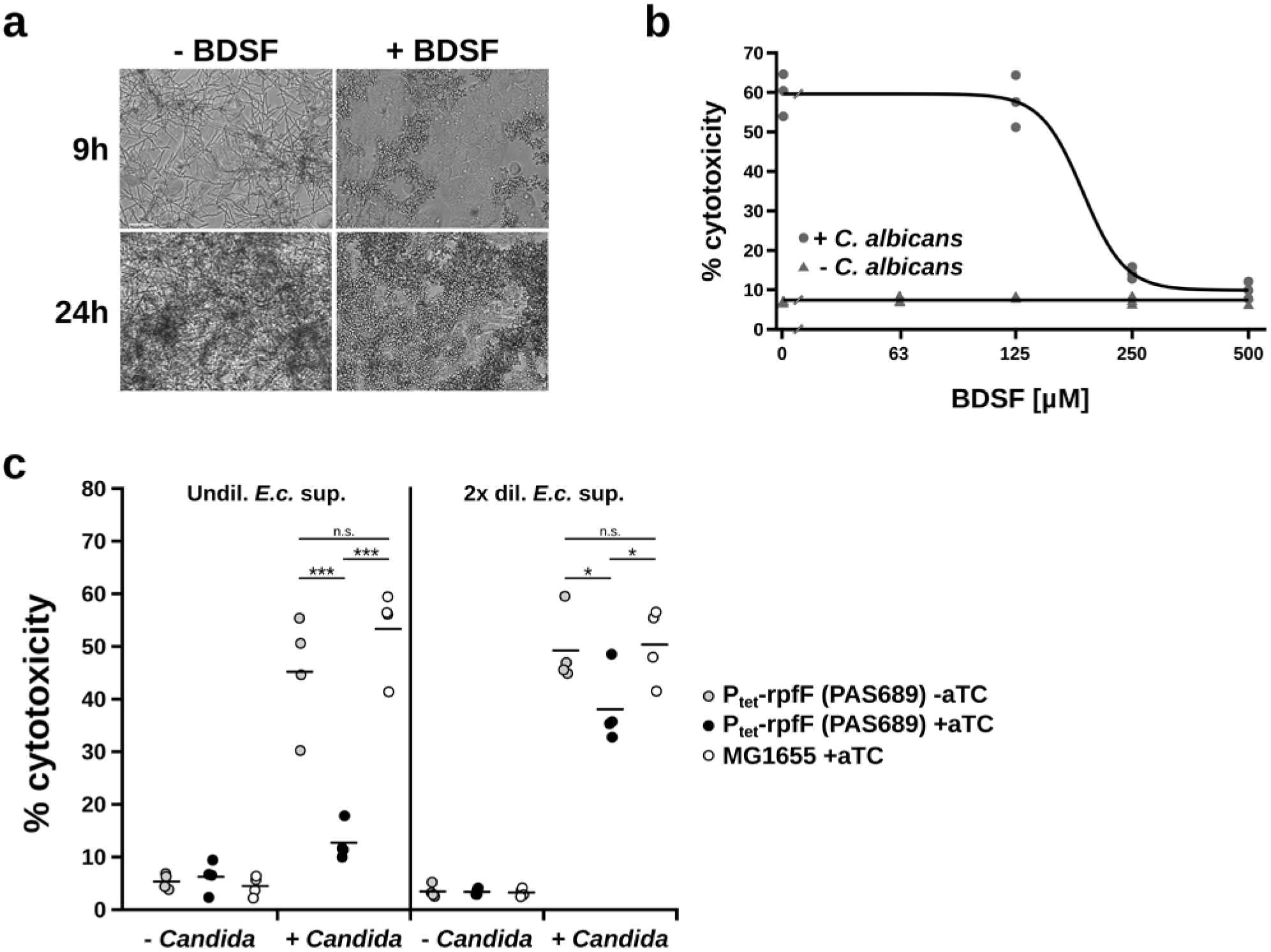
BDSF produced by *E. coli* protects epithelial cells from *C. albicans*-induced damage. (a) Caco-2 epithelial cells were treated with *C. albicans* in the presence or absence of 1 mM BDSF for 9 hours and 24 hours, respectively. Scale bar represents 50 μm. (b) Caco-2 cells were treated with *C. albicans* in the presence of increasing concentrations of BDSF and cytotoxicity was determined after 20 hours by measuring LDH release. A four-parameter log-logistic dose-response model was fitted to the data. (c) Caco-2 cells were treated with *C. albicans* in the presence of *E. coli* supernatants and cytotoxicity was determined after 20 hours. Means of biological replicates are indicated. One-tailed Student’s t-test: *P<0.05; ***P<0.001; n.s.: not significant;

BDSF producing *E. coli* protect epithelial cells from being damaged by *C. albicans*. We treated epithelial cells with *C. albicans* and *E. coli* culture supernatants and measured LDH release after interaction. As expected, in the presence of fungal cells we observed increased cytotoxicity when treated either with wild-type *E. coli* control supernatants from bacteria not carrying *rpfF* or with supernatants from uninduced bacterial cultures (**Figure 4c**). However, treatment with supernatants from aTC-induced PAS689 cells strongly reduced cytotoxicity in a dose-dependent manner. Thus, our data indicate that bacterial cells producing BDSF can protect epithelial cells from *C. albicans*-induced damage.

### Sensing and hypha inhibition by engineered *E. coli*

The *C. albicans* sensing and inhibiting activities can be combined to create a sense-and respond system by coupling the 4-HPA sensor to the *rpfF* gene (**Figure 5a**). To determine if the system can work in commensal strains, the sense-and-respond system was tested in a mouse commensal *E. coli* strain NGF-1^31^. Upon induction with 4-HPA the BDSF concentration in supernatants of cells carrying the sense-and-respond system (PAS694) increased over time (**Figure 5b**). Furthermore, induction with *C. albicans* culture supernatants led to a similar increase in BDSF levels whereas for uninduced cells the inhibitor levels stayed low. To determine if bacteria carrying the sense-and-respond system are able to inhibit hypha formation, we treated *C. albicans* cells under hypha inducing conditions with *E. coli* culture supernatants that had been induced with either 4-HPA or fungal supernatants and determined fungal cell morphology. *E. coli* PAS694 was able to efficiently inhibit hypha formation only upon induction with 4-HPA or fungal supernatants (**Figure 5c**). In contrast, treatment with control supernatants from PAS693 cells lacking RpfF or cells that were not treated with HPA or fungal supernatants allowed extensive filamentation.

**Figure 5.**
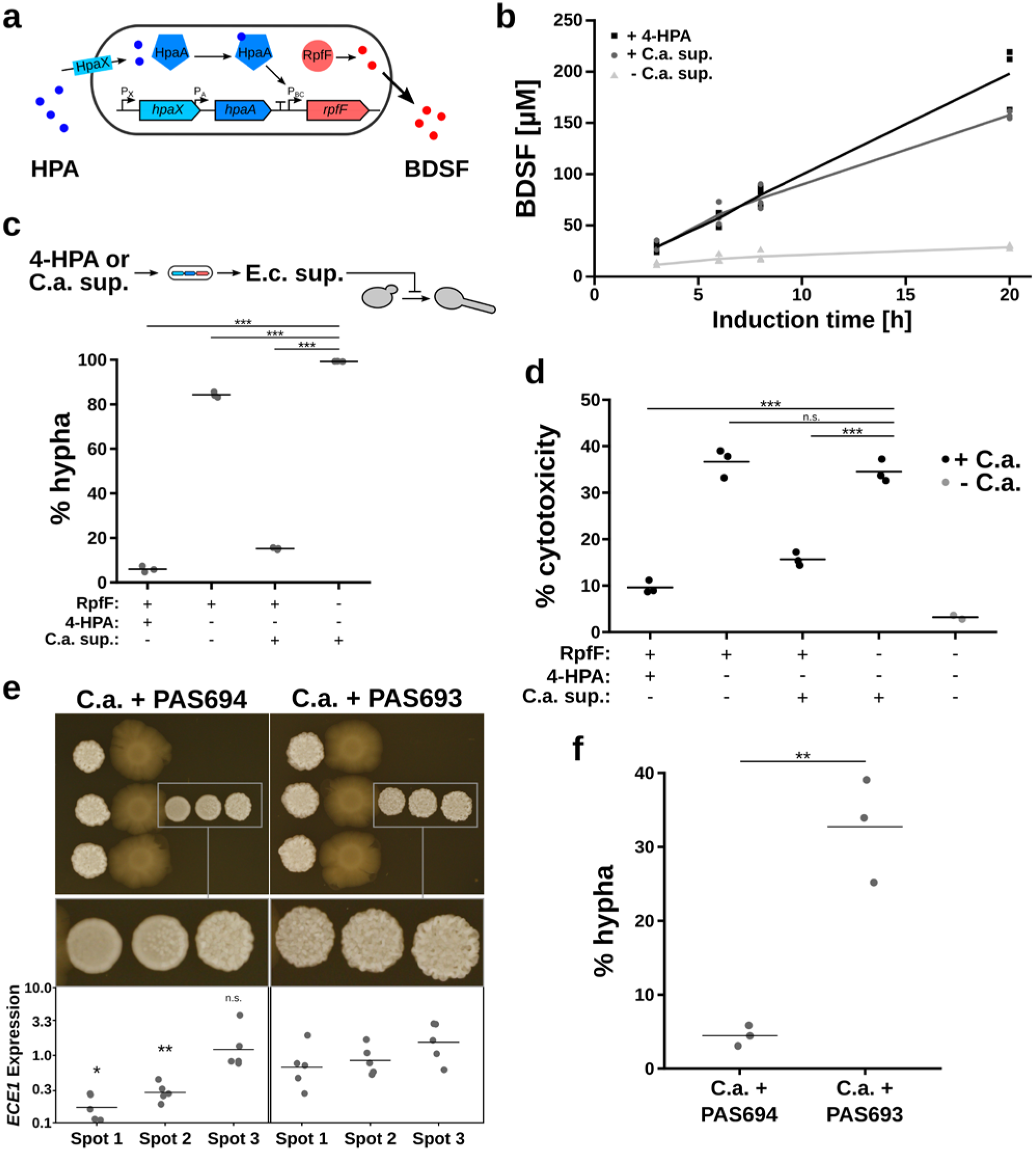
Engineered *E. coli* can sense *C. albicans*, inhibit hypha formation and protect epithelial cells. (a) Design of a *C. albicans* sense-and-respond system. Engineered *E. coli* can import HPA produced by *C. albicans* via the transporter HpaX. HpaA binds to HPA and induces expression of RpfF, which produces the hypha inhibitor BDSF. (b) BDSF levels in supernatants of *E. coli* cells carrying the *C. albicans* sense-and-respond system (PAS694) were determined by HPLC after induction with 100 μM 4-HPA or 0.1x *C. albicans* culture supernatant. (c) *E. coli* PAS694 supernatants harvested 20 hours after induction as described in (b) were incubated with *C. albicans* yeast cells under filament-inducing conditions in minimum medium with glycerol at 37°C and morphology was determined after 3 hours. *E. coli* PAS693 (- RpfF) served as control. (d) *E. coli* PAS694 and PAS693 supernatants prepared as described in (c) were used to treat epithelial cells infected with *C. albicans*. Epithelial damage was determined by measuring LDH release. (e) *C. albicans* and *E. coli* PAS694 were cocultured on agar plates under filament-inducing conditions. Filament inhibition was determined based on colony morphology. *E. coli* PAS693 served as control. *C. albicans* spots were harvested and used for RNA isolation and RT-qPCR analysis of *ECE1* expression levels. Spot 1 is closest to *E. coli*. (f) *C. albicans* and *E. coli* PAS694 and PAS693, respectively, were cocultured on agar plates as described in (e). *C. albicans* spots closest to *E. coli* were harvested and cell morphology was determined microscopically. (b-f) Means of biological replicates are indicated. One-tailed Student’s t-test: *P<0.05; **P<0.01; ***P<0.001; n.s.: not significant relative to PAS693 control;

Bacteria carrying the sense-and-respond system can protect epithelial cells from *C. albicans*-mediated killing. We incubated fungal cells with Caco-2 cell monolayers in the presence or absence of bacterial culture supernatants and determined cytotoxicity by measuring LDH release. A clear reduction in cytotoxicity compared to control supernatants from uninduced cells or cells lacking RpfF was observed only in the presence of supernatants from *E. coli* PAS694 cells, which have been induced by 4-HPA or fungal supernatants prior to harvest (**Figure 5d**). Decreased cytotoxicity and hypha formation in 4-HPA-induced samples compared to samples treated with *C. albicans* supernatants (**Figure 5c and d**) is due to higher BDSF levels upon 4-HPA induction (**Figure 5b**).

*E. coli* cells carrying the sense-and-respond system are able to inhibit hypha formation directly. *C. albicans* growth on agar plates yields an easy readout of filamentation based on colony morphology; smooth colonies indicate the presence of yeast cells whereas wrinkling indicates filamentation^32^. First, fungal cells were spotted on agar plates and incubated for 24 hours to allow the production of HPA. Then, engineered bacteria and subsequently again *C. albicans* were spotted in proximity onto the same agar plate and incubated for additional 48 hours prior to imaging. Only *E. coli* PAS694 carrying the sense-and-respond system were able to inhibit filamentation as indicated by a smooth fungal colony morphology in proximity of the bacterial spots (**Figure 5e**). The inhibiting activity decreased with increased distance to the bacterial cells indicating a gradient of BDSF production by the engineered *E. coli*. Furthermore, we observed no significant inhibition with the PAS693 control strain lacking RpfF.

To determine virulence gene expression, we harvested *C. albicans* cells upon coculture on agar plates and determined the expression levels of the well-known hypha-specific virulence genes *ECE1* (Candidalysin-encoding gene)^8^ and *HWP1* (adhesin)^32^ by RT-qPCR (**Figure 5e** and **Figure S1f**). Only in proximity of *E. coli* cells carrying the sense-and-respond system (C.a. + PAS694 spot 1 and 2) a significant repression of virulence genes was observed. *E. coli* PAS693 served as control. To quantify hypha inhibition, we harvested *C. albicans* cells upon 24 hour coculture on agar plates and determined cell morphology by microscopy. Upon coculture with *E. coli* cells carrying the sense-and-respond system (PAS694) we observed a significant reduction in *C. albicans* hypha formation compared to PAS693 control cells (**Figure 5f**).

Taken together, these data indicate that *E. coli* cells carrying the *C. albicans* sense-and-respond system are able to induce BDSF production in the presence of the fungus thereby inhibiting hypha formation as well as virulence factor expression and protecting epithelial cells from fungal-mediated damage.

## Discussion

In this study, we present the construction of a synthetic sense-and-respond system directed against a human fungal pathogen. We identified HPA as a novel secreted molecule produced by *C. albicans*, one of the most common fungal pathogens infecting humans. Furthermore, we built a sensor able to detect fungal HPA and coupled it to the production of the hypha inhibitor BDSF. Finally, we were able to show that *E. coli* cells carrying the sense-and-respond system can inhibit hypha formation in *C. albicans* and protect epithelial cells from fungal-mediated damage.

To the best of our knowledge, there have been no reports describing the production of HPA by *C. albicans* or any other *Candida* species so far. Similar to tyrosol, HPA production in *C. albicans* is increased in the presence of tyrosine indicating that both molecules might be produced from the same precursors. In *Saccharomyces cerevisiae* HPA and tyrosol are produced via the Ehrlich pathway^33^. *C. albicans* uses the same pathway for production of tyrosol^34^. Thus, HPA might also be produced via the Ehrlich pathway in *C. albicans*. Tyrosol is able to reduce the lag phase in growth upon dilution into minimal medium and it can stimulate hypha induction in *C. albicans*^35^. Given the structural similarity between 4-HPA and tyrosol as well as the possibility of common precursors in biosynthesis the two molecules might have overlapping biological functions. Interestingly, the 4-HPA sensor can be found not only in *E. coli* W strains but also in the human commensal *E. coli* strain HS^36^. Thus, HPA might also have a function in bacterial-fungal communication for example in the human intestine. In fact, 4-HPA has been detected in human fecal samples, although its source remains to be determined^37,38^.

Targeting filamentation of *C. albicans* as an antifungal strategy has gained interest more recently. A number of substances inhibiting the transition from the yeast to the hyphal morphology have been described^39^. Among them, BDSF has been shown to efficiently inhibit hypha formation^13^. We could show that *E. coli* cells expressing RpfF are able to produce the inhibitor at levels sufficient for efficient hypha inhibition. In addition, by engineering fatty acid metabolism, BDSF production could potentially be even further increased^30^. Interestingly, BDSF has also been used to inhibit virulence gene expression in *Pseudomonas aeruginosa*^40^. It inhibits expression of a type III secretion system and has been used to protect epithelial cells as well as zebrafish from *P. aeruginosa*-mediated killing. Thus, our engineered sense-and-respond system could potentially be easily adapted to be effective against *P. aeruginosa* by coupling expression of RpfF to existing sensors for this important bacterial pathogen^24^. Various members of the human microbiota have been shown to produce inhibitors of *C. albicans* hypha formation or virulence^13,14,16^. Often the actual inhibitory molecule is not known and constitutively produced^41–43^. These observations suggest that bacteria inhabiting the gastrointestinal tract along with *C. albicans* may utilize an inhibitory strategy to enhance their ability to co-exist with *C. albicans* in their common niche. Our sense-and-respond system represents the first engineered system able to exploit this strategy.

Filamentation is critical for active penetration of epithelia, which is the main mechanism used by *C. albicans* to invade intestinal epithelial cells^44,45^. Thus, inhibiting hypha formation could avoid *C. albicans* invasion of the intestinal epithelium. However, filamentation is not absolutely required for intestinal colonization since cells lacking the transcription factor Efg1, which are defective in hypha formation, show higher colonization levels in mice compared to wild-type cells at least within 3 days upon gavage (Pierce, 2012)^46^. Thus, our approach would not affect fungal growth and colonization in general, but inhibit epithelial penetration and translocation into the bloodstream.

Our engineered bacterium detects a fungal pathogen and induces production of an inhibitor of *C. albicans* virulence factors. The system has the ability to protect epithelial cells from *C. albicans*-mediated damage. Engineered probiotics could lower treatment costs since the active substance is produced by the microbe^17^. Furthermore, inhibiting virulence factors instead of growth can lower the risk of resistance development^7,17^. Thus, our approach represents a novel strategy to prevent fungal infections with the potential to reduce treatment costs and counteract the development of antifungal resistance.

## Methods

### Bacterial and fungal strains and growth conditions

Bacterial and fungal strains used in this study are listed in Table S1. *E. coli* strains were routinely grown in LB medium at 37°C. *C. albicans* was routinely grown in YPD medium at 30°C. Minimum medium as described in Boon *et al*.^13^ was used for sensor induction with *C. albicans* supernatants and coculture experiments and was supplemented with 0.4% glycerol or *N*-acetylglucosamine when indicated. M9 medium supplemented with 0.1% casamino acids and 0.4% glycerol was used for dose-response experiments with 4-HPA and tyrosol. For solid media 1.5% agar was added for LB plates and 2% agar for YPD plates. Antibiotics were used at the following concentrations: 100 μg/mL spectinomycin, 25 μg/mL chloramphenicol, 50 μg/mL kanamycin and 100 μg/mL ampicillin.

### Mammalian cell culture

Caco-2BBE human colorectal carcinoma-derived (Caco-2) intestinal epithelial cells (obtained from Harvard Digestive Disease Center) were routinely cultured in Dulbecco’s Modified Eagle Medium (DMEM) containing 4.5 g/L glucose (ATCC) supplemented with 10% fetal bovine serum (FBS), 100 units/mL penicillin and 100 μg/mL streptomycin (Pen/Strep) at 37°C in a humidified incubator under 5% CO_2_ in air.

### Plasmid and strain construction

Plasmids and oligonucleotides used in this study are listed in Tables S2 and S3, respectively. *E. coli* NEB5α was used for cloning. For the construction of pTet a *tetR-P_tetA_* fragment was amplified from the PAS132 genome with primers tet_XmaJI_fw & tet_BamHI_re and BamHI/XmaJI cloned into the pBAD His A backbone amplified with primers pBAD_fw & pBAD_XmaJI_re. Then, pTet-GFP was constructed by inserting the gfpmut2 gene from pUA66^47^ into pTet via BamHI/PstI. The *hpaXA-P_BC_* fragment was amplified from the ATCC11105 genome with primers hpaX_XmaJI_fw & PhpaBC_XbaI_re and XmaJI/XbaI cloned into pTet-GFP replacing the *tetR-P_tetA_* part to obtain pHpaXA-GFP. For the construction of pCDF-HpaXA-GFP a *hpaXA*-GFP fragment was amplified from pHpaXA-GFP with primers hpaX_XmaJI_fw & pHpaXA_ApaI_re and XmaJI/ApaI cloned into the pCDFDuet-1 backbone amplified with primers pACYC_ApaI_fw & pACYC_XmaJI_re. The *hpaX* gene was removed from pCDF-HpaXA-GFP by amplification with primers hpaA_XmaJI_fw & pACYC_XmaJI_re and religation upon XmaJI digest to obtain pCDF-HpaA-GFP. The *B. cenocepacia rpfF* gene with an N-terminal 6xHis-tag was obtained as gBlock from Integrated DNA Technologies. For the construction of pCDF-HpaXA-rpfF the 6xHis-*rpfF* fragment was amplified from the gBlock with primers Bcam0581_oe_fw & Bcam0581_oe_re and cloned into the pCDF-HpaXA-GFP backbone amplified with primers PBC_oe_re & pCDF_oe_fw to replace the *gfp* gene. Plasmids were transformed into different *E. coli* strains via heat shock.

To obtain pKD3-Ptet-rpfF the *tetR-P_tetA_* fragment was amplified from the PAS132 genome with primers tet_fw & tet_re, the pKD3 vector backbone was amplified with primers pKD3_fw & pKD3_tetR_re and both fragments were combined with the 6xHis-*rpfF* gBlock using Gibson assembly. For the construction of PAS689 the Cm^R^-*tetR-P_tetA_-rpfF* fragment was amplified from pKD3-Ptet-rpfF with primers pKD3int_fw & pKD3int_re and integrated into the *araB-araC* locus of *E. coli* TB10 as described^48^. The locus was then transferred to *E. coli* MG1655 by P1*vir* transduction as described^31^. The streptomycin-resistance mutation *rpsL* (*lys42arg*) was transferred into *E. coli* NGF-1 as described^31^.

### Tyrosol and HPA measurement by HPLC-MS

*C. albicans* SC5314 supernatants were prepared by diluting overnight cultures to an OD_600_ of 0.25 in minimum medium. For tyrosine supplementation 1 mM tyrosine was added to the medium. For hypha induction the medium included 4 g/L *N*-acetylglucosamine. Fungal supernatants were harvested by centrifugation after incubation at 30°C for yeast cultures and 37°C for hyphal cultures for 24 hours with shaking and sterile filtered using 0.22 μm filters. Samples were acidified by addition of 100 mM H_2_SO_4_ and extracted using Sep-Pak Light C18 cartridges (Waters). Samples were reduced in a speedvac benchtop concentrator at 50°C prior to HPLC-MS analysis.

HPLC-MS analysis of standards and extracts was carried out using an Agilent 1260 Infinity HPLC system equipped with an Agilent Eclipse Plus C18 column (100 × 4.6 mm, particle size 3.5 μm, flow rate: 0.3 mL/min, solvent A: dd.H_2_O/0.1% (v/v) NH_4_OH, solvent B: acetonitrile, injection volume: 7.5 μL) connected to an Agilent 6530 Accurate-Mass Q-TOF instrument. The following gradient was used for separation and analysis (time/min, %B): 0, 0; 13, 100; 18, 100; 19, 0, 25, 0. The mass spectrometer was operated in negative ESI mode and the autosampler was kept at 4°C. MS-fragmentation experiments were carried out using collision-induced dissociation (CID) fragmentation.

To quantitate tyrosol and HPA levels in extracts, a standard curve was recorded using freshly prepared tyrosol and HPA standards dissolved in dd.H2O/acetonitrile (85/15 (v/v)). Samples were kept on ice or at 4°C. After HPLC-MS analysis, extracted ion current (EIC) peaks (m/z (tyrosol): 137.05-137.07, m/z (HPA): 151.0-151.05) were automatically integrated using the MassHunter Workstation Software (version: B.07.00). A plot of peak area versus concentration was used to generate a linear fit and quantitate levels found in extracts.

### BDSF measurement by HPLC

*E. coli* PAS694 and PAS693 overnight cultures were diluted 1:100 in minimum medium with 0.4% glycerol and spectinomycin and incubated at 37°C for 4 hours. Glycerol supplementation was done to enhance *E. coli* growth. For induction 100 μM 4-HPA or 0.1x *C. albicans* supernatant in minimum medium with tyrosine prepared as described above was added and cultures were incubated at 37°C with shaking. Supernatants were harvested at different time points and sterile filtered using 0.22 μm filters. BDSF extraction was done essentially as described^49^ with some modifications. First, 200 μL culture supernatant were acidified by addition of 0.3 M HCl and extraction was done twice with 200 μL ethyl acetate. Extracts were dried in a speedvac benchtop concentrator at 40°C and dissolved in 100 μL methanol. Five μL were used for analysis on an Agilent 1200 HPLC system using a Zorbax SB-C18 column (3.0 x 150 mm, 3.5 μm). The mobile phase was 80% methanol with 0.1% formic acid and the flow rate was 0.4 mL/min. BDSF was detected using a diode array detector set to 220 nm (4 nm band width). Quantification was done using BDSF standards spiked into medium and extracted as described above.

### Dose-response analysis

For dose-response analysis with 4-HPA and tyrosol as inducers, *E. coli* overnight cultures were diluted 1:100 in M9 medium containing different concentrations of 4-HPA or tyrosol and incubated at 37°C for 20 hours with shaking. GFP levels and OD_595_ were determined using a Victor3V plate reader (PerkinElmer). After blank correction GFP levels were normalized for OD_595_ to obtain RFU values. A four-parameter log-logistic dose-response model was fitted to the data using R version 3.2.3 and the *drc* package.

*C. albicans* culture supernatants for dose-reponse analysis were prepared as described above. *E. coli* overnight cultures were diluted 1:100 in minimum medium supplemented with 0.4% glycerol. Fungal culture supernatants were added at different dilutions and cultures were incubated at 37°C for 20 hours with shaking. GFP and OD_595_ measurement as well as RFU calculation was done as described above.

### Bacterial-fungal coculture

To determine sensor induction on agar plates, *C. albicans* SC5314 overnight cultures were washed once and diluted in PBS buffer to OD_600_ of 0.1 and 1 μL was spotted on LB agar plates supplemented with 0.4% glycerol to support fungal growth and spectinomycin. Plates were incubated for 24 hours at 37°C. *E. coli* PAS691 and PAS697 overnight cultures were diluted to OD_600_ of 0.1 in PBS buffer and 1 μL spotted next to *C. albicans*. Plates were imaged after 24 hours incubation at 37°C. Images were taken using an imaging station equipped with a Canon EOS Rebel T3i camera and GFP excitation/emission filters.

To determine hypha inhibition on agar plates, *C. albicans* SC5314 suspensions were spotted on YPGlcNacSpec plates containing 10 g/L yeast extract, 20 g/L peptone and 4 g/L *N-*acetylglucosamine allowing hypha formation. Cell suspensions were prepared as described above. Plates were incubated for 24 hours at 37°C. Then *E. coli* PAS694 and PAS693 suspensions were spotted next to *C. albicans* as described above and plates were further incubated at 37°C for 6 hours. Finally, *C. albicans* suspensions were spotted in different distances to the *E. coli* spots and plates were incubated at 37°C for 48 hours prior to imaging as described above. Quantification of hyphal cells was done by resuspending *C. albicans* spots after 24 hours coculture in 0.5 ml PBS containing 0.05% Tween 20. Cells were sonicated for 10 seconds using a tip sonicator to disperse clumps prior to microscopy using an inverted Zeiss Axiovert 200M microscope. At least 800 cells were counted per sample.

For bacterial-fungal cocultures in liquid medium *C. albicans* SC5314 overnight cultures were washed twice and diluted in minimum medium with spectinomycin to OD_600_ of 0.12. *E. coli* PAS691 overnight cultures were diluted to OD_600_ of 0.0063 in minimum medium in the absence or presence of *C. albicans*. Cells were incubated in CellView glass bottom culture dishes at 37°C and images were taken using an inverted Nikon Ti2 fluorescence microscope over time in 15 minute intervals.

### Hypha inhibition assay

*C. albicans* SC5314 overnight cultures were washed once and diluted to OD_600_ of 0.2 in PBS buffer. Five μL cell suspension were added to 100 μL DMEM medium with 5% FBS and different concentrations of BDSF in clear bottom 96 well plates. Cells were incubated at 37°C for 3 hours to induce hypha formation prior to imaging on an inverted Nikon TE2000 microscope to determine cell morphology. More than 50 cells were counted for each condition and the experiment was repeated 3 times. The percentage of cells showing hyphal morphology was determined. A four-parameter log-logistic dose-response model was fitted to the data using R version 3.2.3 and the drc package.

For hypha inhibition by *E. coli* supernatants upon aTC induction PAS689 and MG1655 overnight cultures were diluted to OD_600_ of 0.05 in DMEM medium without phenol red and Pen/Strep in the absence or presence of 100 ng/mL aTC and incubated at 37°C with shaking. Supernatants were harvested at different time points, sterile filtered using 0.22 μm filters and 100 μL aliquots transferred to clear bottom 96 well plates. Upon addition of *C. albicans* as described above and 5% FBS to induce hypha formation cells were incubated at 37°C for 3 hours. Imaging and analysis was done as described above.

For hypha inhibition by *E. coli* supernatants upon 4-HPA or *C. albicans* induction PAS694 and PAS693 overnight cultures were diluted 1:100 in minimum medium with glycerol and spectinomycin and incubated at 37°C for 4 hours. For induction 100 μM 4-HPA or 0.1x *C. albicans* supernatant in minimum medium with tyrosine prepared as described above was added and cultures were incubated for 20 hours. Supernatants were harvested, sterile filtered and 50 μL aliquots transferred to clear bottom 96 well plates. Upon addition of *C. albicans* as described above and 50 μL 2x minimum medium with 8 g/L *N*-acetylglucosamine to induce hypha formation cells were incubated at 37°C for 3 hours. Imaging and analysis was done as described above.

### Analysis of virulence gene expression

*C. albicans* CA58 overnight cultures were washed once and resuspended at OD_600_ of 0.25 in DMEM medium or aTC-induced PAS689 and MG1655 supernatants prepared as described above and 5% FBS were added to induce hypha formation. For BDSF treated samples the inhibitor concentration was 1 mM. Samples were incubated at 37°C with shaking for 3 hours prior to imaging on an inverted Nikon TE2000 fluorescence microscope.

For analysis of *ECE1* and *HWP1* expression by RT-qPCR fungal cells were resuspended in 1 ml PBS and harvested by centrifugation. RNA extraction, DNAse treatment and qPCR analysis was done as described previously^50^. *PAT1* and *RIP1* reference genes were used for normalization^51^.

### Fungal-mammalian coculture and cytotoxicity assay

Caco-2 cells were seeded at 5 x 10^4^ cells/well in 96 well plates and incubated for 3 days to form a monolayer. *C. albicans* SC5314 overnight cultures were diluted to OD_600_ of 0.1 in YPD medium and incubated at 30°C until they reached OD_600_ 1.0. Cells were harvested, washed twice in PBS buffer and resuspended in DMEM medium. To determine inhibition of cytotoxicity by BDSF the Caco-2 medium was changed to DMEM 5% FBS without phenol red containing different concentrations of BDSF. Then, 1 x 10^4^ *C. albicans* yeast cells per well were added and samples were incubated at 37°C 5% CO_2_ for 20 hours. Finally, LDH release into supernatants was determined using the CytoTox 96 Non-Radioactive Cytotoxicity Assay (Promega) according to the manufacturer’s instructions. To determine inhibition of cytotoxicity by PAS689, aTC-induced culture supernatants were prepared as described above for hypha inhibition assays. Caco-2 medium was replaced by *E. coli* supernatants supplemented with 5% FBS. After addition of *C. albicans* yeast cells samples were incubated and LDH release determined as described above.

To determine inhibition of cytotoxicity by PAS694, 4-HPA-induced and *C. albicans*-induced bacterial supernatants were prepared as described above for hypha inhibition assays. PAS693 supernatants served as controls. Supernatants were diluted 4x in DMEM 10% FBS medium without phenol red and added to Caco-2 cells. After addition of *C. albicans* yeast cells samples were incubated and LDH release determined as described above.

For imaging of Caco-2 cells treated with *C. albicans* in the absence and presence of BDSF, samples were prepared as described above. Images were taken after 9 hours and 24 hours interaction, respectively, using an inverted Leica DM IL LED microscope.

### Transwell culture conditions and epithelial barrier measurements

Caco-2 cells were seeded at 1 x 10^5^ cells/cm^2^ in collagen-coated transwell inserts (Corning) containing 200 μL DMEM medium in the apical compartment and 800 μL in the basal compartment. Cells were incubated at 37°C 5% CO_2_ for 18-20 days and the medium was changed every 2-3 days. *C. albicans* SC5314 cells were prepared as described above for cytotoxicity assays and 1 x 10^5^ cells were added to the apical compartment in the absence or presence of 500 μM BDSF. The medium in both compartments was removed after 12 and 24 hours. Fresh DMEM was added to the basal compartment and DMEM with BDSF was used for the apical compartment. Transepithelial electrical resistance and permeability measurements were used to determine epithelial barrier function exactly as described^52^.

## Acknowledgments

We are grateful to Bernhard Hube for providing *C. albicans* strain CA58 and Julia Köhler for providing *C. albicans* SC5314. This work was funded in part by Defense Advanced Research Projects Agency Grant HR0011-15-C-0094 and funds from the Wyss Institute for Biologically Inspired Engineering. M.T. was supported by an Erwin Schrödinger Fellowship (J3835) from the Austrian Science Fund. T.W.G. was supported by a Leopoldina Research Fellowship (LPDS 2014-05) from the German National Academy of Sciences. C.A.K and L.M. were supported by NIH grant AI118898.

## Author contributions

M.T. and P.A.S. designed the project. M.T., T.W.G. and L.M. performed experimental work. M.T. and T.W.G. analyzed the results. C.A.K. gave technical and conceptual advice. M.T. and P.A.S. wrote the manuscript and T.W.G. and C.A.K. edited the manuscript.

## Supplementary Information

**Figure S1.**
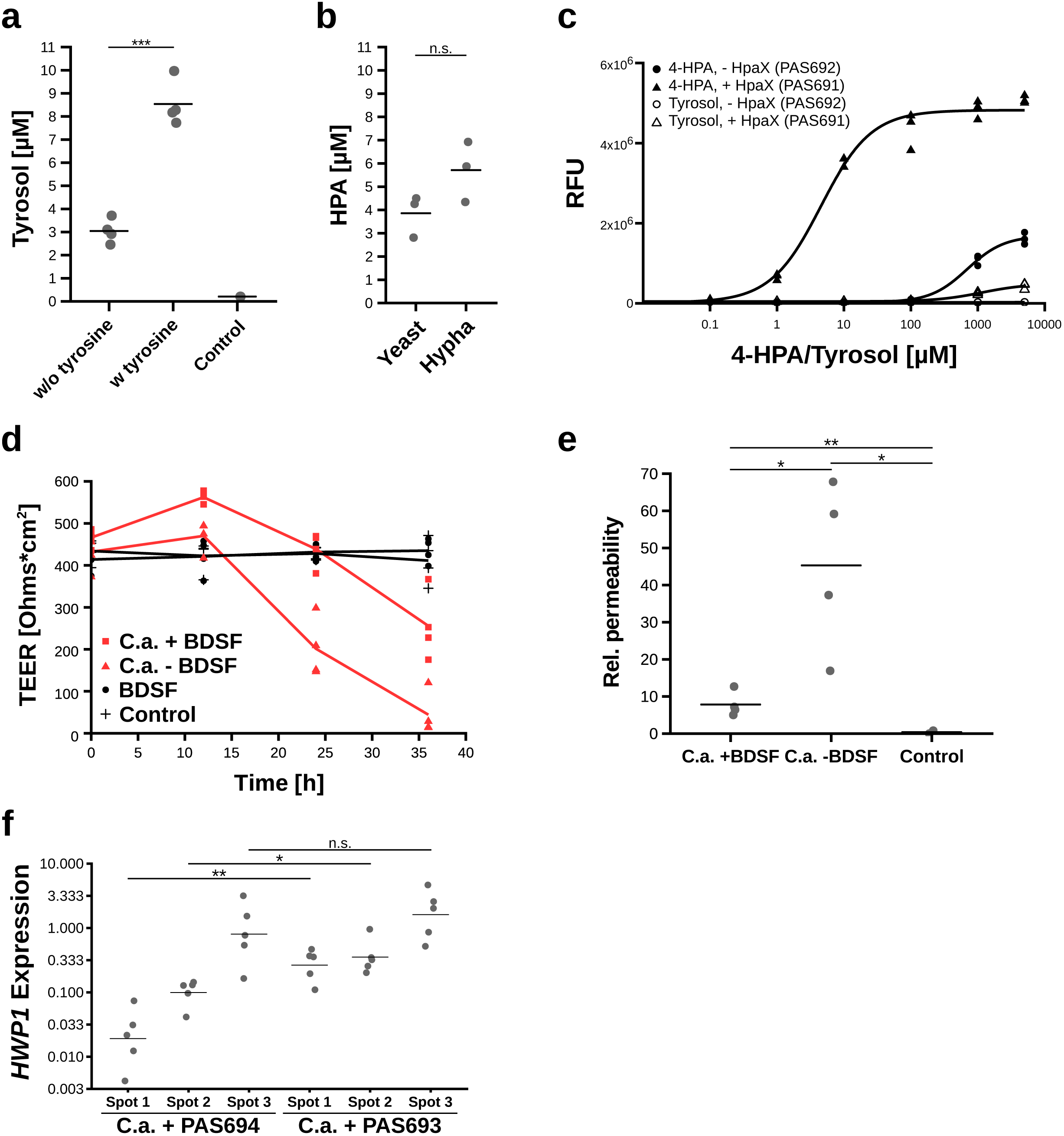
(a) Tyrosol levels in *C. albicans* culture supernatants were determined by HPLC-MS. Addition of tyrosine increases production. (b) HPA levels in *C. albicans* yeast and hyphal culture supernatants were determined by HPLC-MS. (c) Dose-response of an *E. coli* 4-HPA sensor to 4-HPA and tyrosol. GFP was placed under the control of a 4-HPA inducible promoter, which by HpaA. Expression of the 4-HPA transporter HpaX increases sensitivity. A four-parameter log-logistic dose-response was fitted to the data. (d) Caco-2 cells cultured in transwell plates were treated with *C. albicans* -/+ 500 μM BDSF in the compartment and transepithelial electrical resistance (TEER) was measured at the indicated time points. (e) Caco-2 cells treated as described in (d) and epithelial permeability was determined after 36 hours interaction. (f) *C. albicans* and *E. coli* and PAS693, repestively, were cocultured on agar plates under hypha-inducing conditions as described in Figure 5e. *C.* spots were harvested and used for RNA isolation and RT-qPCR analysis of *HWP1* expression levels. Spot 1 is closest to *E. coli*. (a-f) Means of biological replicates are indicated. One-tailed Student’s *t*-test: \**P*<0.05; \*\**P*<0.01; n.s.: not significant;

**Table S1:**
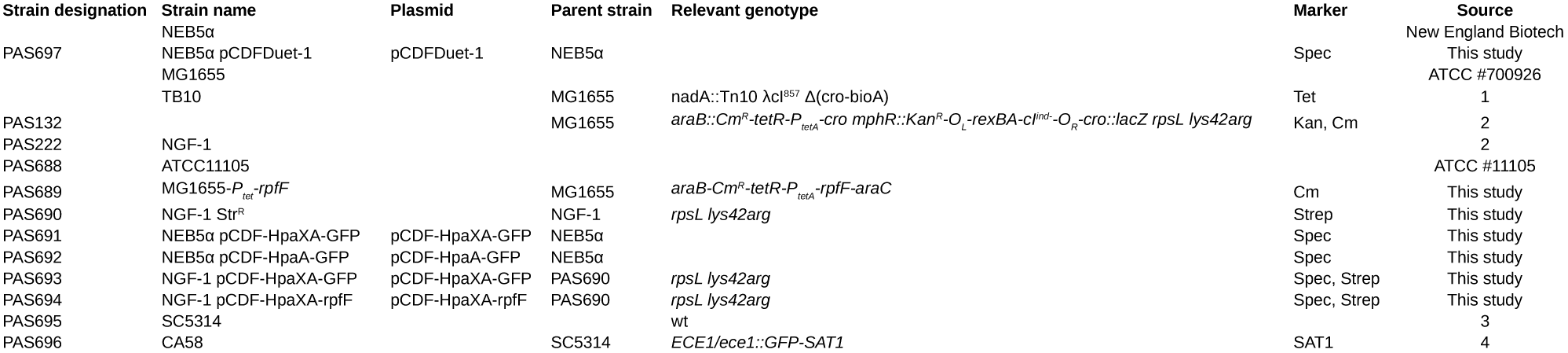
Bacterial and fungal strains used in this study. Strain designation Strain name.

**Table S2:**
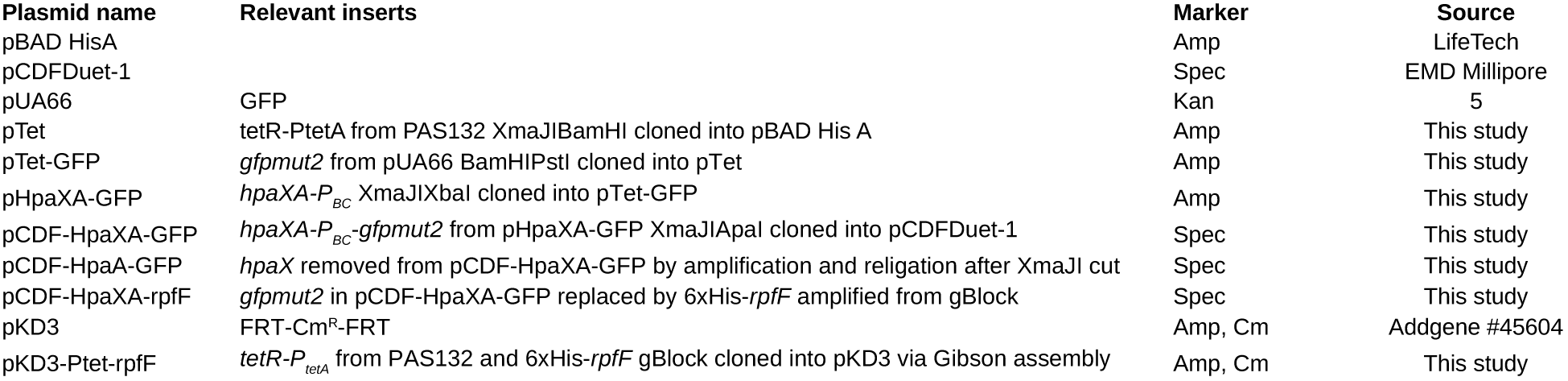
Plasmids used in this study. Plasmid name Relevant inserts.

| Plasmid name | Relevant inserts | Marker | Source |
| --- | --- | --- | --- |
| pBAD HisA |  | Amp | LifeTech |
| pCDFDuet-1 |  | Spec | EMD Millipore |
| pUA66 | GFP | Kan | 5 |
| pTet | tetR-PtetA from PAS132 XmaJIBamHI cloned into pBAD His A | Amp | This study |
| pTet-GFP | *gfpmut2* from pUA66 BamHIPstI cloned into pTet | Amp | This study |
| pHpaXA-GFP | *hpaXA-P<sub>BC</sub>* XmaJIXbaI cloned into pTet-GFP | Amp | This study |
| pCDF-HpaXA-GFP | *hpaXA-P<sub>BC</sub>-gfpmut2* from pHpaXA-GFP XmaJIApaI cloned into pCDFDuet-1 | Spec | This study |
| pCDF-HpaA-GFP | *hpaX* removed from pCDF-HpaXA-GFP by amplification and religation after XmaJI cut | Spec | This study |
| pCDF-HpaXA-rpfF | *gfpmut2* in pCDF-HpaXA-GFP replaced by 6xHis-*rpfF* amplified from gBlock | Spec | This study |
| pKD3 | FRT-Cm<sup>R</sup>-FRT | Amp, Cm | Addgene #45604 |
| pKD3-Ptet-rpfF | *tetR-P<sub>tetA</sub>* from PAS132 and 6xHis-*rpfF* gBlock cloned into pKD3 via Gibson assembly | Amp, Cm | This study |

**Table S3:**
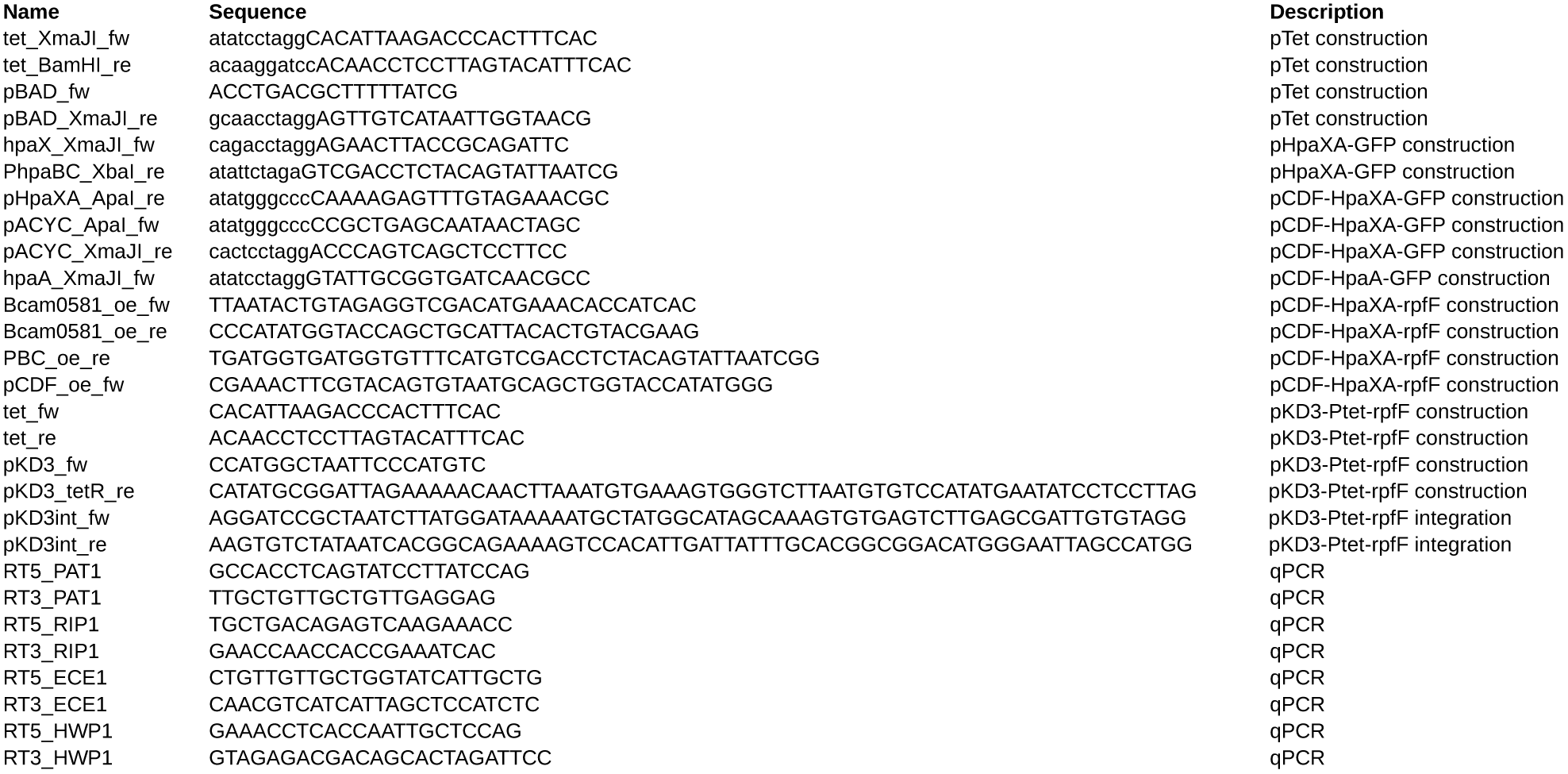
Oligonucleotides used in this study. Name.

